# Interpretable and Robust Machine Learning for Exploring and Classifying Soundscape Data

**DOI:** 10.1101/2024.11.07.622465

**Authors:** Arpit Omprakash, Rohini Balakrishnan, Robert Ewers, Sarab Sethi

## Abstract

The adoption of machine learning in Passive Acoustic Monitoring (PAM) has improved prediction accuracy for tasks like species-specific call detection and habitat quality estimation. However, these models often lack interpretability, and PAM generates vast amounts of non-informative data, as soundscapes are typically information sparse.

Here, we developed ecologically interpretable methods that accurately predict land use from audio while filtering unwanted data. Audio from habitats in Southern India (evergreen forests, deciduous forests, scrublands, grasslands) was collected and categorised by land use (reference, disturbed, and agriculture). We used Gaussian Mixture Models (GMMs) on top of a Convolutional Neural Network (CNN)-based feature extractor to predict land use.

Thresholding based on likelihood values from GMMs improved model accuracy by excluding uninformative data, enabling our method to outperform models such as Random Forests and Support Vector Machines. By analysing areas of acoustic feature space driving predictions, we identified “keystone” soundscape elements for each land use, including both biotic and anthropogenic sources.

Our approach provides a novel method for ecologically meaningful interpretation and exploration of large acoustic datasets independent of specific feature extractors. Our study paves the way for soundscape monitoring to deliver robust and trustworthy habitat assessments on scales that would not otherwise be possible.

## Introduction

Curbing the ongoing biodiversity crisis calls for fast and accurate biodiversity and habitat quality assessments to guide the development and implementation of appropriate conservation methods (IUCN, 2024; Katswera et al., 2022). Traditional biodiversity and habitat quality monitoring methods, such as field surveys, are tedious, expensive, and often limited to a few specific sites at any time (Moore, 2009). Furthermore, many survey methods, such as mark-recapture and mist netting, are invasive and affect surveyed animal populations (Moore, 2009; Zemanova & Zemanova, 2020). Passive Acoustic Monitoring (PAM) offers an easily scalable, inexpensive, and non-invasive alternative to traditional biodiversity surveys (Heinicke et al., 2015; Rosenthal & Ryan, 2000).

Vocalising biodiversity in an area depends on factors such as habitat type, quality and anthropogenic activity, and affects a region’s soundscape (Pijanowski et al., 2011). A soundscape is the amalgamation of all sounds emanating from a landscape (Pijanowski et al., 2011). The soundscape contains information not only on the biodiversity of a given region but also on the habitat quality and other environmental and anthropogenic variables such as land use intensity (Müller et al., 2022; Tucker et al., 2014; Turlington et al., 2024). One of the fundamental goals of ecoacoustics is to study the relationship between habitat quality and soundscapes and whether there are unifying acoustic features of habitat quality across ecosystems that can be assessed using PAM (Sánchez-Giraldo et al., 2021; Sangermano, 2022; Shamon et al., 2021).

The desire to analyse soundscapes for habitat level insights has led to the development of ecoacoustic indices and soundscape-based machine-learning models to estimate species richness, activity, and habitat quality (Buxton et al., 2018; Sethi et al., 2022; Shamon et al., 2021). Many studies employ ecoacoustic indices as proxies for ecosystem health and habitat quality (Krause & Farina, 2016; Sánchez-Giraldo et al., 2021). However, these indices are often sensitive to non-biological sounds (geophony and anthropophony) and fail to work in complex environments such as tropical forests and urban habitats (Eldridge et al., 2018; Fairbrass et al., 2017; Jorge et al., 2018; Ross et al., 2021). Soundscape-based machine-learning models infer latent features (often called embeddings) from audio data using machine-learning algorithms that can be used for downstream tasks like audio segmentation and classification (Sethi et al., 2020; Zaugg et al., 2023). Although these models show greater accuracy in predicting metrics of habitat quality, such as above-ground biomass, across a broader range of ecosystems compared to ecoacoustic indices (Sethi et al., 2020), they are still fallible (Sethi et al., 2023) and there is no simple method of determining how these models produce predictions, hindering interpretability.

Most machine learning models used for PAM are based on Convolutional Neural Network (CNN) architectures. These models use a spectrogram of the audio file as input to compute latent features called “acoustic feature embeddings” (or just ‘embeddings’) for different audio files (Krizhevsky et al., 2012). The success of these models over other ecoacoustic indices indicates that these embeddings contain more relevant information than single or multi-modal acoustic indices (McGinn et al., 2023; Sethi et al., 2022). However, these embeddings are non-semantic, abstract numerical representations of audio data, with no method to ascertain the nature of information contained in these embeddings. Many of these models might draw accurate conclusions from biased latent features, like highly correlated background noise (Gibb et al., 2024; Nieto-Mora et al., 2024; Ribeiro et al., 2016). With the ever-increasing size of acoustic datasets, it becomes difficult to regulate and understand the mechanisms by which these models make classifications. Thus, these models might give higher prediction accuracies for a given task, even while misinterpreting the task at hand. This fundamentally decreases the trustworthiness of these model predictions, especially on previously unseen data. Recently, the non-interpretability of these successful models has been recognised as a barrier to the advancement of the scientific study of soundscapes, especially when the fundamental relationships between soundscape patterns and landscape dynamics, ecosystem structure, and function are still not fully understood (Gibb et al., 2024; Nieto-Mora et al., 2024).

Another common issue in most PAM studies is the failure to recognize the sparseness of informational content in soundscape recordings, which arises from the inherently sparse and temporally segregated nature of biophonic sounds (Francomano et al., 2020; Gottesman et al., 2020). Unless sampling for a particular taxon or using a specific recording schedule, the amount of uninformative data collected and fed through any analysis pipeline is enormous.

With the rapid adoption of cheap Automated Recording Units (ARUs), researchers are recording soundscapes across day and night, generating extremely large volumes of audio data (Gamillo, 2023; Roe et al., 2021; Sethi et al., 2021). Current methods to identify and isolate information-rich parts from soundscape audio data are practically nonexistent in ecoacoustics. At the same time, techniques such as foreground-background element separation are prevalent in other fields of acoustic scene classification, indicating the need for such algorithms in ecoacoustics (Devalraju & Rajan, 2022; Dhanunjaya et al., 2020). Soundscape informational content is sparse, and there are not a lot of tools that can help filter these non-informational pieces of audio data. Developing a method to sort through this massive amount of ecoacoustic data generated from long-term monitoring projects would help us train better models and devise better temporal sampling protocols for habitat quality estimation.

The overarching question we address in this study is whether we can develop an interpretable, soundscape-based model to predict land use disturbance across different ecosystems using audio alone. A binary classifier to distinguish between soundscapes from two extreme land use classes (e.g., forest vs agriculture) provides a simple starting point that verges on the trivial. However, we tackle the more difficult task of challenging the algorithm to classify intermediate levels of habitat disturbance (e.g., managed forest) that form a gradient between these two extremes. Moreover, existing soundscape methods seldom provide ecological insight and a large portion of audio data that is collected will likely be uninformative: (1) we do not know which soundscape elements can be used to distinguish between different land uses; and (2) we do not know whether there are unifying acoustic elements that can be used to classify land use gradients across different ecosystems. Here, we extend our method to extract exactly this type of information from unstructured soundscape recordings, thereby providing more detailed insight into the signature acoustic elements that define habitat disturbance gradients.

This study presents a novel density estimation-based approach to ecoacoustic data analysis that harnesses the proven predictive power of CNN-derived acoustic embeddings whilst delivering interpretable and explainable decisions. First, we demonstrate the framework to make land use classifications and show that using thresholding on the probabilistic predictions of the model, we can eradicate uninformative soundscape components and improve model accuracy. Secondly, we show how we can use the same framework to identify, isolate, and visualise ecologically relevant soundscape components important in distinguishing different land uses across different ecosystems and habitats, demonstrating its use as a powerful data exploration tool and providing interpretability and credibility to the predictions made by our model.

## Materials and Methods

### Study Sites

We collected data from multiple habitat types: deciduous and evergreen forests, scrublands, and grasslands in Karnataka, India. We identified three broad land use types for each habitat based on the intensity of anthropogenic activity (number of people passing through, forest produce collection, grazing intensity) in different habitats. All land use types in a habitat were geographically close to each other to keep factors other than anthropogenic disturbance, such as climatic conditions, constant. The land use types can be classified into three broad categories – Reference (plots with minimal anthropogenic disturbance), Disturbed (plots with considerable and regular anthropogenic activities), and Agriculture (agricultural sites including fields and plantations). We used 15 plots (points of recording) to collect data in each habitat, five in each land use type (except in the Evergreen habitat, where we used four plots in each land use type due to space constraints). Each plot was at least 200 m apart from all other plots and the land use type’s boundary to prevent auditory interference across plots and minimise edge effects. A few of the deployed recorders malfunctioned, slightly decreasing the number of plots sampled for audio across habitats (total number of plots sampled across all habitats = 53). A table containing detailed information regarding the various sampling sites and land-use plots is given in the supplementary section (see Table T1).

**Figure 1.**
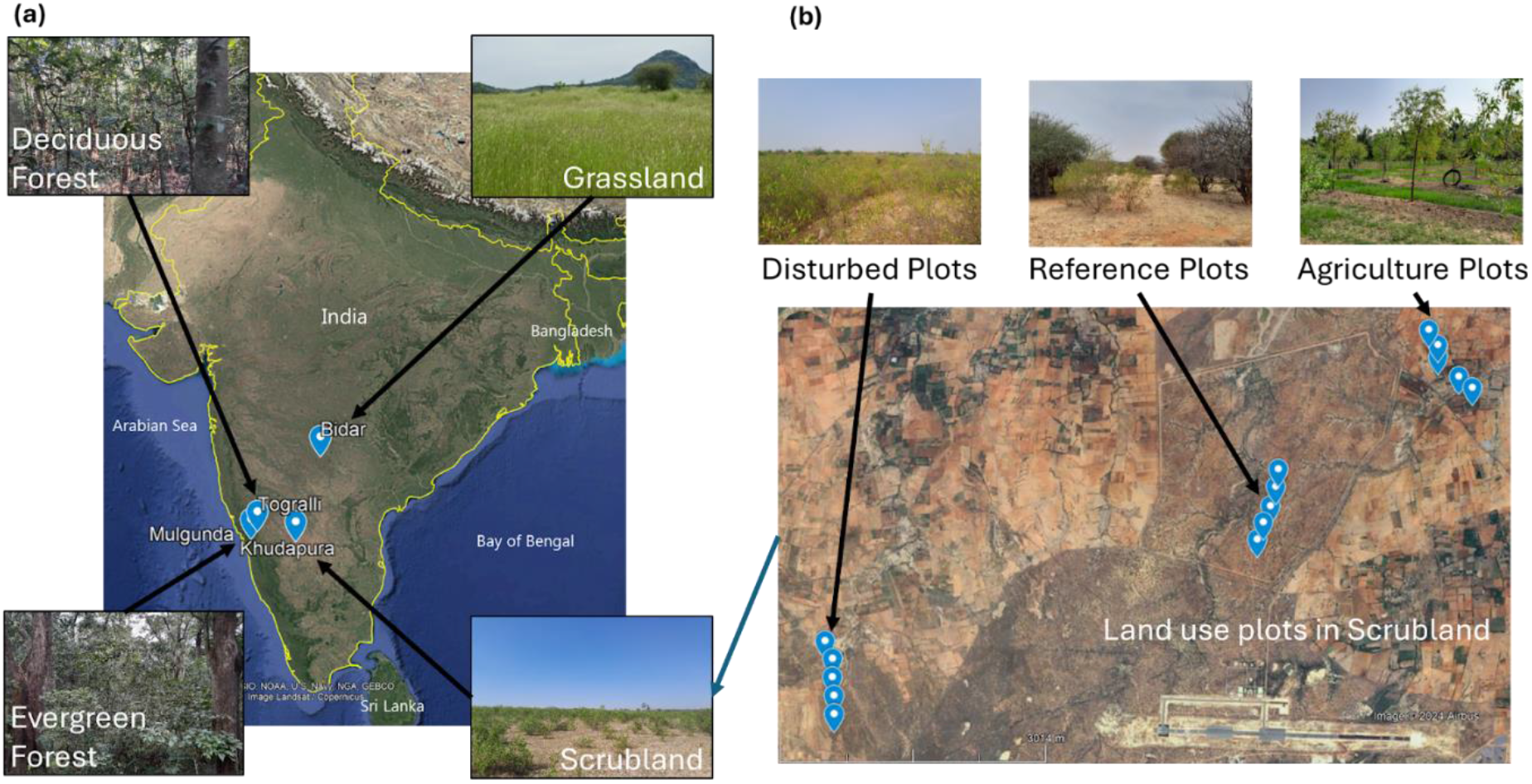
Study sites and study design. (a) Location of the study sites with respect to the Indian subcontinent. Sample images taken from all four habitats are shown. (b) An enlarged view of the scrubland habitat shows the various plots in each land use, along with sample images from all three land uses. Five plots in each land use type were set up to record audio in each habitat.

**Figure 2.**
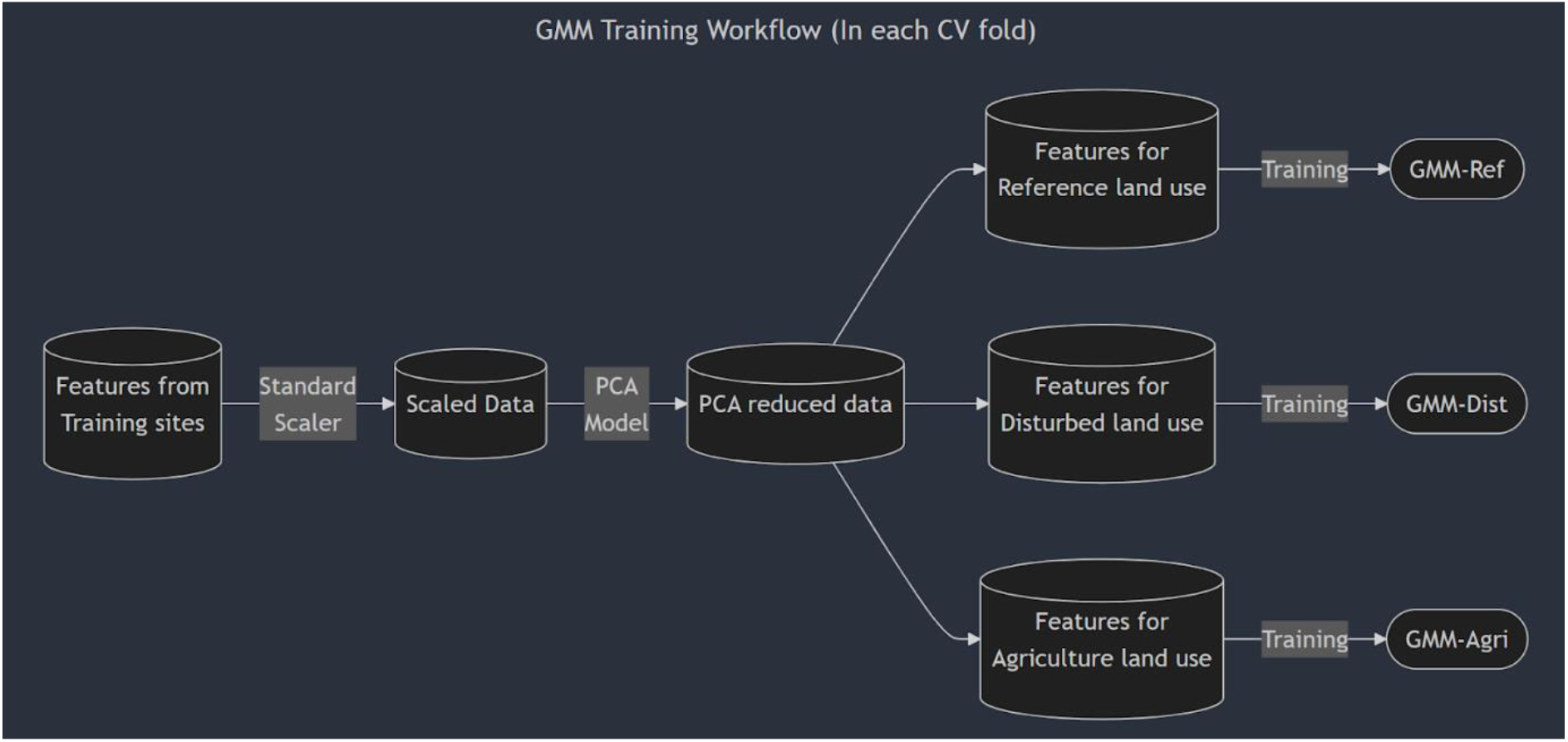
GMM Training Workflow in a given fold of cross-validation. We first standardised the columns of the audio feature vector. It was followed by a PCA to reduce the dimensions of the training data. This PCA-reduced data was then segregated based on the land use, and three GMMs were trained for the three land uses in each habitat.

### Audio Data Collection and Feature Extraction

Audio data were collected in all the plots using AudioMoths v1.2.0 (Hill et al., 2018). These automatic recording devices were placed on tree branches at a height of approximately 2 m above ground. They were programmed to record at a sampling rate of 192 kHz for five consecutive days in each plot at a 50 % duty cycle, i.e., capturing alternative 5-minute audio segments. Since there were fewer plots for evergreen habitat (n = 12), we used a slightly higher duty cycle (66.67 %). The recordings in each habitat (across all the land use types) were made simultaneously. All audio data from the various study sites were collected between January and March 2023. In total, 3176 hours and 22 min of audio data were collected across all the habitats (Table T1).

Each audio file was divided into chunks of 0.96 s. These chunks were then resampled at 16 kHz using a Kaiser window. Using this resampled audio, log-scaled Mel-frequency spectrograms were computed (96 temporal frames, 64 frequency bands). Each spectrogram was passed through the VGGish CNN from Google’s Audioset project (Gemmeke et al., 2017; Hershey et al., 2017), generating a 128-dimensional feature embedding of the audio sample. Based on the land use of the plot in which the audio was recorded, each audio sample (0.96 s chunk of audio) was assigned to a given land use category (Reference, Disturbed, or Agriculture). This constituted the primary dataset for all the following analyses.

### Training the Gaussian Mixture Model Classifier

We used a Bayesian Gaussian Mixture Model with 100 components and diagonal covariance matrices to build the classifier for our framework (see supplementary). Unless otherwise specified, any further mention of GMM should be assumed to be a Bayesian GMM. We carried out four-fold cross-validation using the StratifiedGroupKFold method of sklearn (Pedregosa et al., 2011). StratifiedGroupKFold creates a stratified split of the data in each fold of the cross-validation, ensuring that the proportion of various land uses remains similar in the training and testing splits while also ensuring that the data from a given site is restricted to either the training or testing dataset, thus preventing data leakage. The training data in each fold consisted of audio feature vectors and corresponding land use labels from 12 randomly selected plots (four plots per land use), and the testing data consisted of the remaining three plots (one per land use). The training audio feature vectors were further processed in each cross-validation fold before training models. The feature vectors were first standardised using the StandardScaler method of sklearn (Pedregosa et al., 2011), and a PCA (Principal Components Analysis) was carried out to reduce the number of dimensions while preserving 95% variation in the training data of the current fold. The PCA-reduced data points were then segregated based on land use, and three GMMs were trained, one corresponding to each land use. A schematic of the training process is shown below:

### Predicting using the Gaussian Mixture Model Classifier

#### Generating baseline predictions

Audio feature vectors from the testing data were taken to generate predictions for the baseline model. In each cross-validation fold, the feature vectors were first transformed using the corresponding scaler and PCA models (from the model training phase) for the fold to complete the data pre-processing. The PCA-reduced test data were then passed through all three GMMs for the cross-validation fold to generate log-likelihood scores for each land use model, e.g., S_Ref_, S_Dist_, S_Agri_. Based on these scores, likelihood ratios were calculated for each model using the following formulae:

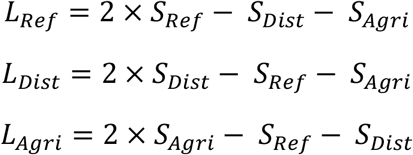

The likelihood ratio quantifies the likelihood of observing the test data point under the different models. This formulation was adapted from the two-model format of the likelihood ratio or Wilks test to work for three competing statistical models (Wilks, 1938). A high likelihood ratio value for reference indicates a higher probability of observing the data under the GMM-Ref model, and so on. We also compared models using the log-likelihood scores and the likelihood ratios to make predictions and test if our likelihood ratio formulation works. In all cases, the model’s accuracy using the likelihood ratios was either equal to or greater than the model’s accuracy using the log-likelihood scores. Thus, we used the likelihood ratio to generate model predictions. By comparing the likelihood ratios for the different models, we could select the model under which the data is most likely to be observed to make a prediction. For a given audio embedding X, the land use was predicted according to the following equation:

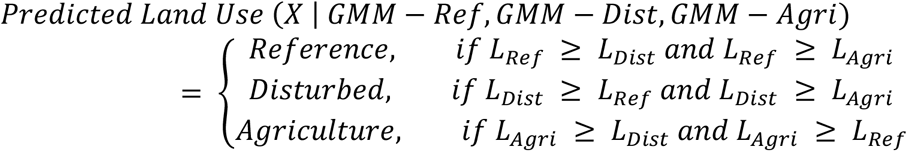

#### Generating thresholded predictions

We introduced thresholding in the classifier to drop uninformative data points. We used the training data to generate the thresholds, ensuring no data leakage. We first made predictions on the training data and extracted the likelihood ratios for all the models, i.e., GMM-Ref, GMM-Dist, and GMM-Agri, for all true positive predictions. The threshold was then set as a percentile-based cut-off based on the distribution of these likelihood ratios. A threshold of 30 indicates that after generating the distribution of likelihood ratios for the three GMMs, we identified the value at the 30^th^ percentile in each distribution (it might vary for the three models), say Ref_30_, Dist_30_, and Agri_30,_ which are then treated as the actual cut-off values for their respective GMM models. Any likelihood ratio value less than Ref_30_ was discarded for GMM-Ref; any likelihood ratio value less than Dist_30_ was discarded for GMM-Dist, and the same was true for GMM-Agri and Agri_30._ The following equation summarises the final likelihood ratio calculation for the thresholded model:

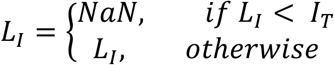

Where,

L_I_ is the likelihood ratio for the model I (I ∈{Ref, Dist, Agri})

I_T_ is the threshold value for model I at the percentile T threshold

When making predictions on the test dataset, we passed the audio embeddings into the three models for the corresponding cross-validation fold. Once the likelihood ratios were obtained, they were thresholded. The final land use label prediction for a given feature embedding X at threshold T was then made as follows:

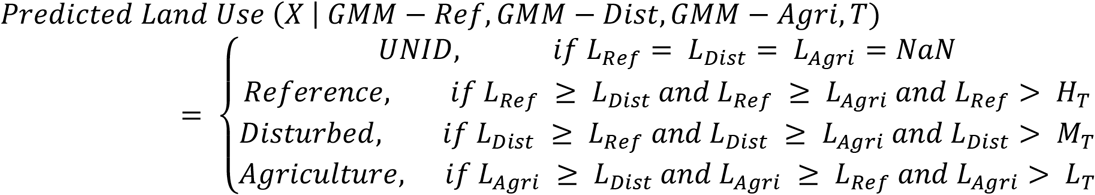

For each habitat, models were run at various percentile thresholds ranging from 1 to 99 in increments of 3. A model F1 score gradient vs threshold plot was made (see supplementary Figure S1). The optimal percentile threshold for a given habitat was determined by the first inflexion point of the curve (the point where the slope of the curve first becomes zero). This represented the first local maxima for the F1 score.

Thresholding and changes in predictions

We encountered 3 cases when thresholding model predictions:

1. No change: If all three likelihood ratios were below the threshold, there was no change in the baseline model prediction post-thresholding.
2. Change in Predictions: If the likelihood ratio for the baseline prediction was below the threshold and at least one other likelihood ratio was above the threshold, the model predictions changed.
3. Dropping data points: If all three likelihood ratios were below the threshold, the data point was dropped from the model predictions.

Dropping data points always led to a decrease in either true positive or false positive predictions. In contrast, a change in prediction might lead to an increase or decrease in true positive and false positive predictions. In our analyses, the number of dropped data points far outweighed the number of prediction changes at sufficiently high threshold values. Thus, at high thresholds, it was safely assumed that any decrease in true or false positives was driven by dropping data points. At the same time, any increase was attributed to changes in predictions.

### Estimating model accuracy

The predictions generated in each cross-validation fold were aggregated, and accuracy metrics, such as precision, recall, and F1 score, were calculated based on the aggregated prediction set. Since the data were evenly distributed among all three land use classes for each habitat, all accuracy metrics calculated were macro averaged, i.e., the metrics were calculated for each class or land use, and then the unweighted mean was calculated and represented as the final value of the metric.

### Comparison with other classifiers

To benchmark the prediction accuracy of the GMM model, we also trained and evaluated Support Vector Classifier (SVC) and Random Forest Classifier (RFC) models following the same stratified group four-fold cross-validation methods and training using the same acoustic feature vector dataset. We then estimated the precision scores for all the models to make comparisons between the models.

### Visualising the acoustic feature space using GMMs

To visualise the acoustic feature space, audio embeddings from a single habitat (across land uses) were aggregated, standardised, and transformed by fitting a PCA to reduce dimensions while preserving 95% of the variation. The data was then segregated based on land use, and three GMMs were trained, one for each land use. These models generated for each habitat are referred to as the aggregated model for habitats in later sections. For 2-dimensional visualisation, another PCA was carried out to reduce the dimensions for each transformed audio embedding to two dimensions. The embeddings were grouped into various cells based on their location in the 2-D space. In each cell, the intensity of different colours was based on the average log-likelihood scores for the three models (GMM-Ref, GMM-Dist, GMM-Agri). This method generated acoustic feature space for the baseline and optimally thresholded models.

### Sampling and Labelling audio features

An aggregated model was trained using a given habitat’s data to identify soundscape elements used by the GMM model for land use classification. Land use predictions were made using the aggregated model for all audio embeddings in the habitat at the optimal threshold following the above-mentioned methods. Once the predictions were generated, all the UNID (unidentified) data points were purged from the predicted dataset as the UNID data points were uninformative for land use prediction. GMM components with at least 30 samples were first filtered for all three land uses, and the proportion of true positives was quantified for these components. Five components with the highest proportion of true positives were isolated for each land use GMM. Using the predict_proba method of the GMM in sklearn, the affinity of each data point to a particular GMM component was quantified. Based on the distribution of these affinity scores for a given component, data points with an affinity greater than five percentile for the five selected GMM components were filtered. Thirty data points were randomly sampled for each of the five selected GMM components from the filtered list of data points, giving rise to 450 data points for each habitat to label.

Audio streams of 0.96 s corresponding to each data point were identified, and a buffer region of 1 second on either side of the region of interest was used to extract audio samples corresponding to the best-performing components for each land use across different habitats. These audio samples were then listened to, and the spectrogram visualised (at settings matching that of VGGish, i.e., Sample rate of 16000, STFT window length of 0.025 s, and STFT window hop length of 0.01 s) using Raven Pro, and soundscape elements were identified. Their proportions were quantified to decide on broad labels for the various GMM components.

## Results

### Visualising acoustic feature densities reveals acoustic overlap between land uses

Using a 2-dimensional PCA, we visualised the VGGish embeddings for each habitat to determine how soundscapes from the different land uses were segregated in the audio embedding space. Even in two dimensions, we could identify regions of the acoustic space corresponding to the different land uses and patterns across habitats. Along the main diagonal of the deciduous forest plot (Figure 3a), the acoustic embeddings change from mainly agriculture (left and top parts of the plot) to disturbed to reference (bottom right parts of the plot), indicating that the soundscape components present in audio recordings from reference land use were very different from components present in agricultural land use, with minimal overlap. In contrast, disturbed land use showed some characteristics of both as it lay between both agriculture and reference land uses in the acoustic space and had overlapping regions with both. Across habitats, more areas were coloured orange (a mix of yellow and red) than green (a mix of yellow and blue), indicating more overlaps between acoustic feature embeddings of agriculture and disturbed land uses compared to agriculture and reference land use (Figure 3).

**Figure 3.**
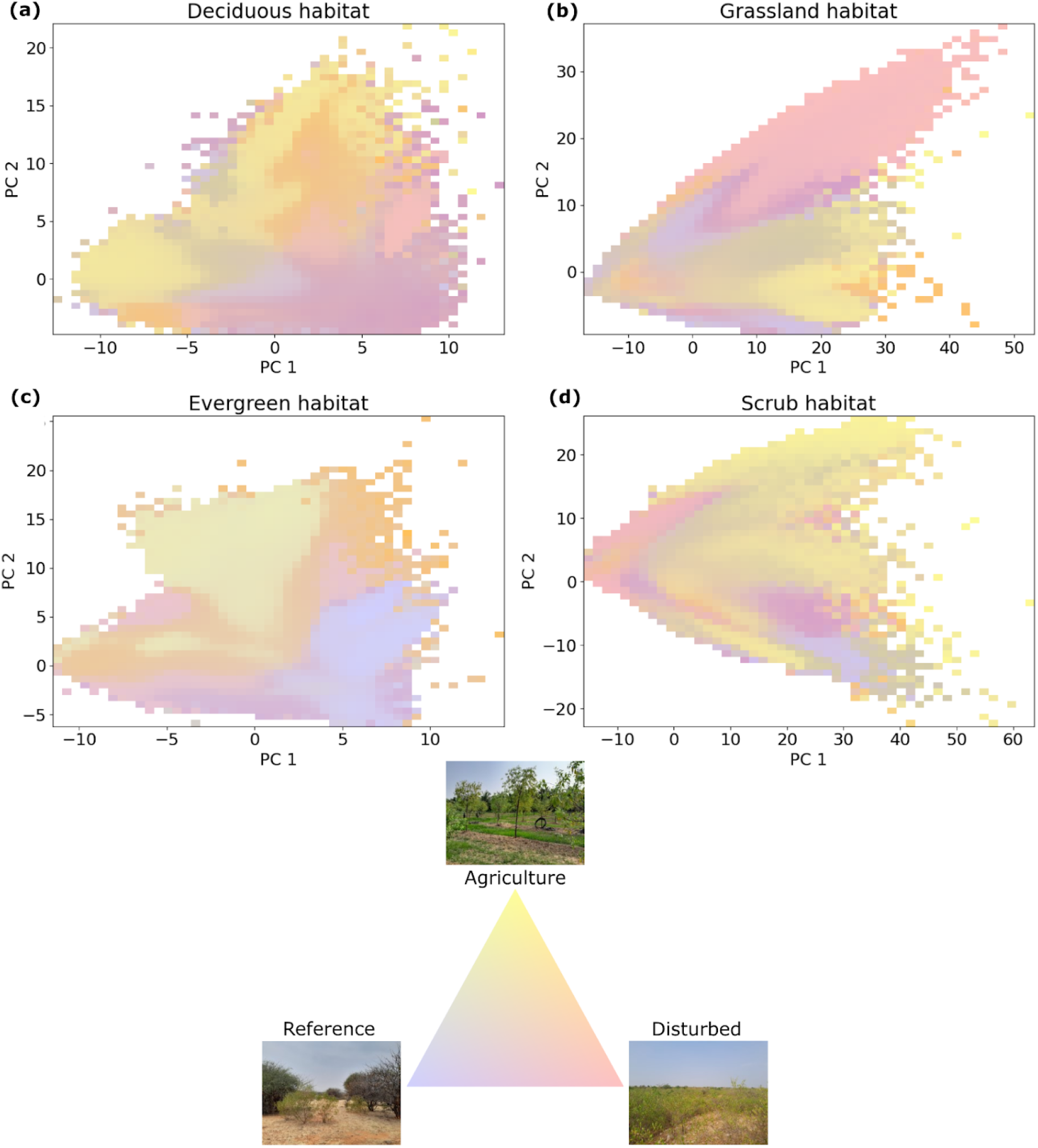
Visualising acoustic features from the different land uses across the four habitats. In each subplot, blue indicates reference habitat, red indicates disturbed habitat, and yellow indicates agricultural land use. Regions belonging to distinct land uses in various habitats can be visually distinguished in the plots. In deciduous habitat, the different land uses separate along the main diagonal, with embeddings from agriculture concentrated in the top-left of the plot and reference audio embeddings concentrated at the bottom-right. Across habitats, there are more overlaps between disturbed and agriculture (demarcated by orange) compared to reference and agriculture (demarcated by green colour), indicating more similarities between soundscapes of disturbed and agriculture when compared to reference and agriculture. This also indicates that the model can separate the acoustic features present in various land uses, and distinct acoustic features and regions in the embedding space correspond to particular land uses.

### Filtering data by thresholding GMM likelihood ratios increases model accuracy across all habitats

We evaluated model performance pre- and post-thresholding at varying threshold levels to understand the effects of thresholding on model accuracy. The classification model without any thresholding (baseline) had precision values of 0.53, 0.48, 0.47, and 0.41 for deciduous forests, evergreen forests, grassland, and scrub habitats, respectively. Increasing the threshold increased model precision, recall, and F1 scores across most habitats (Figure 4 e-h, Figure S2). In the case of scrub habitat, the recall and F1 scores tend to decrease slightly after the 20th percentile threshold (Figure S2). There was also a corresponding increase in the percentage of data points labelled as “UNID” with increasing threshold, indicating a trade-off between model accuracy and the percentage of data points dropped, with more data dropping leading to higher model accuracy (Figure 4 e-h). On average, across habitats, the increase in precision at the optimal threshold was about 12%. The most significant improvement was seen in the case of grassland, where the model precision rose by 20% after thresholding. The scrub habitat had the most negligible improvement, where the precision rose by 2% post-thresholding. Although the increase in the scrub case was marginal, in all the cases, there was an increase in model precision values post-thresholding.

**Figure 4.**
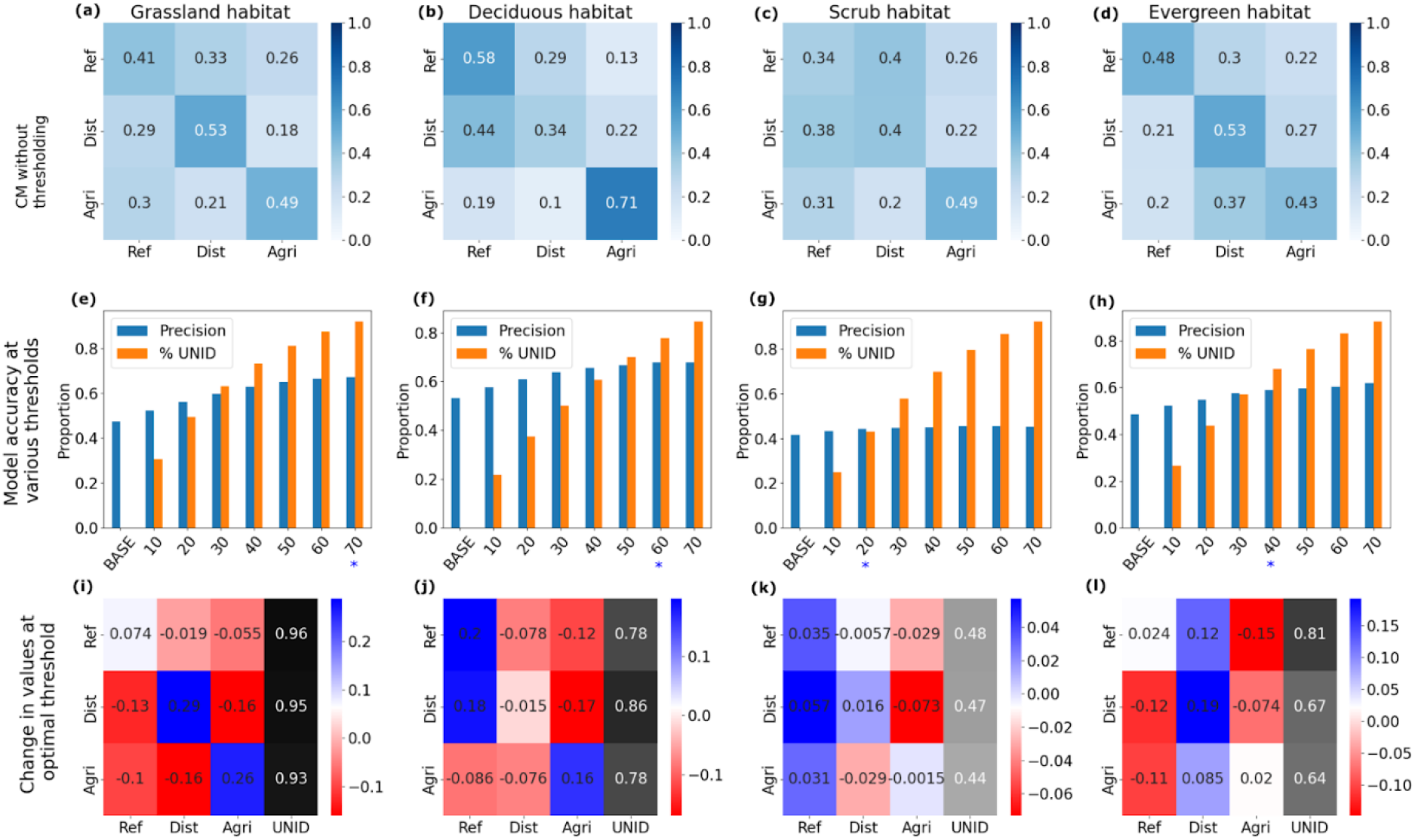
Effect of thresholding on model accuracy. (a-d) Confusion matrices for the baseline model. (e-h) Model precision and percentage of unlabelled data points (UNID) for baseline (BASE) and various thresholds. Increasing the threshold increased model precision and the percentage of UNID data points. Blue stars represent threshold values closest to the optimal threshold for each habitat. (i-l) Differential confusion matrices for all habitats. The differential confusion matrix shows the difference in values for the optimal thresholded model compared to the baseline model. The pattern for grassland habitat best represents how the thresholding increases correct classifications (blue diagonal) and decreases the number of misclassifications (red in all other cells). Similar improvements were seen for the deciduous habitat. In Evergreen, decreasing false positives is the primary driver of improved model performance. Thresholding leads to only marginal improvement in model performance in scrub.

The baseline GMM precision values were either lower or nearly comparable to those of other classification algorithms we tested - Support Vector Classifiers (SVCs) and Random Forest Classifiers (RFCs) (Table 1). However, after optimal thresholding, the precision of GMM classifiers was higher than RFC and SVC across all habitats, indicating the importance of dropping uninformative data points (Table 1). We also find that the model accuracy across different habitats fluctuated similarly for all the models (except SVC in deciduous habitat), with deciduous having the highest precision scores for the baseline (0.53) and RFC models (0.61). In contrast, scrub had the poorest performance across most models (the highest precision achieved was 0.44 using thresholded GMMs). Corroboration with other classification models suggests that the information within soundscapes differed across habitats, and the patterns in performance that we observed were not only caused by our methodological choices.

**Table 1.** Precision scores for various classification models across different habitats.

| Habitat | RFC | SVC | GMM Baseline | GMM Optimal Threshold |
| --- | --- | --- | --- | --- |
| Deciduous | 0.61 | 0.3 | 0.53 | **0.68** |
| Evergreen | 0.53 | 0.43 | 0.48 | **0.59** |
| Grassland | 0.48 | 0.42 | 0.47 | **0.67** |
| Scrub | 0.39 | 0.31 | 0.41 | **0.44** |

### Data filtering by thresholding leads to a drop in uninformative data points across habitats

The effect of thresholding on model predictions was estimated by calculating the change in confusion matrix values for the optimally thresholded model compared to the baseline model. Optimal thresholds for deciduous, evergreen, grassland, and scrub were determined to be 64, 43, 76, and 22 percentiles, respectively, leading to the drop of at least 45% data in all four cases. These thresholds were considered high since they dropped nearly half of the uninformative data points. Thus, any increase in value in the differential confusion matrix (blue colour) was attributed to prediction changes due to thresholding and any decrease (red colour) was attributed to dropping data points (Figure 4 i-l).

For grassland, true positives increased, while false positives decreased post-thresholding (Figure 4i). This indicated that the model dropped most false positive predictions and modified some false positive predictions to true positives, thus increasing model precision, exemplifying the ideal mechanism of increased model accuracy due to thresholding. A similar pattern was observed in deciduous habitats. There were a few misclassifications of disturbed land use as reference, but it was outweighed by the increase in classification accuracy provided by the other two land use classes (Figure 4j). For both evergreen and scrub habitats, there were far more misclassifications in the disturbed and reference columns, respectively. The off-diagonal blue vertical lines in Figures 4k and 4l illustrate that the model misclassified data points from other land uses as disturbed or reference in these habitats, respectively. Similar to the deciduous habitat, the decrease in accuracy due to these misclassifications in the evergreen habitat was overcome by dropping more false positive predictions, thus leading to a decent overall increase in precision (Figure 4l). However, in the case of scrub, only marginal improvement was seen.

The proportion of data points dropped across various land uses in a given habitat was more or less constant with two exceptions: disturbed for deciduous forest and reference for evergreen forest habitats (Figure 4j, 4l). After thresholding, these land uses were also the most misclassified for their respective habitats. In disturbed land use of deciduous forest habitat, the model picked out samples containing silence, indicating a lack of distinguishing soundscape elements (Figure 5). Combined with the higher proportion of unlabelled samples, this might indicate that the model is trying to remove these data points and often misclassifies them.

**Figure 5.**
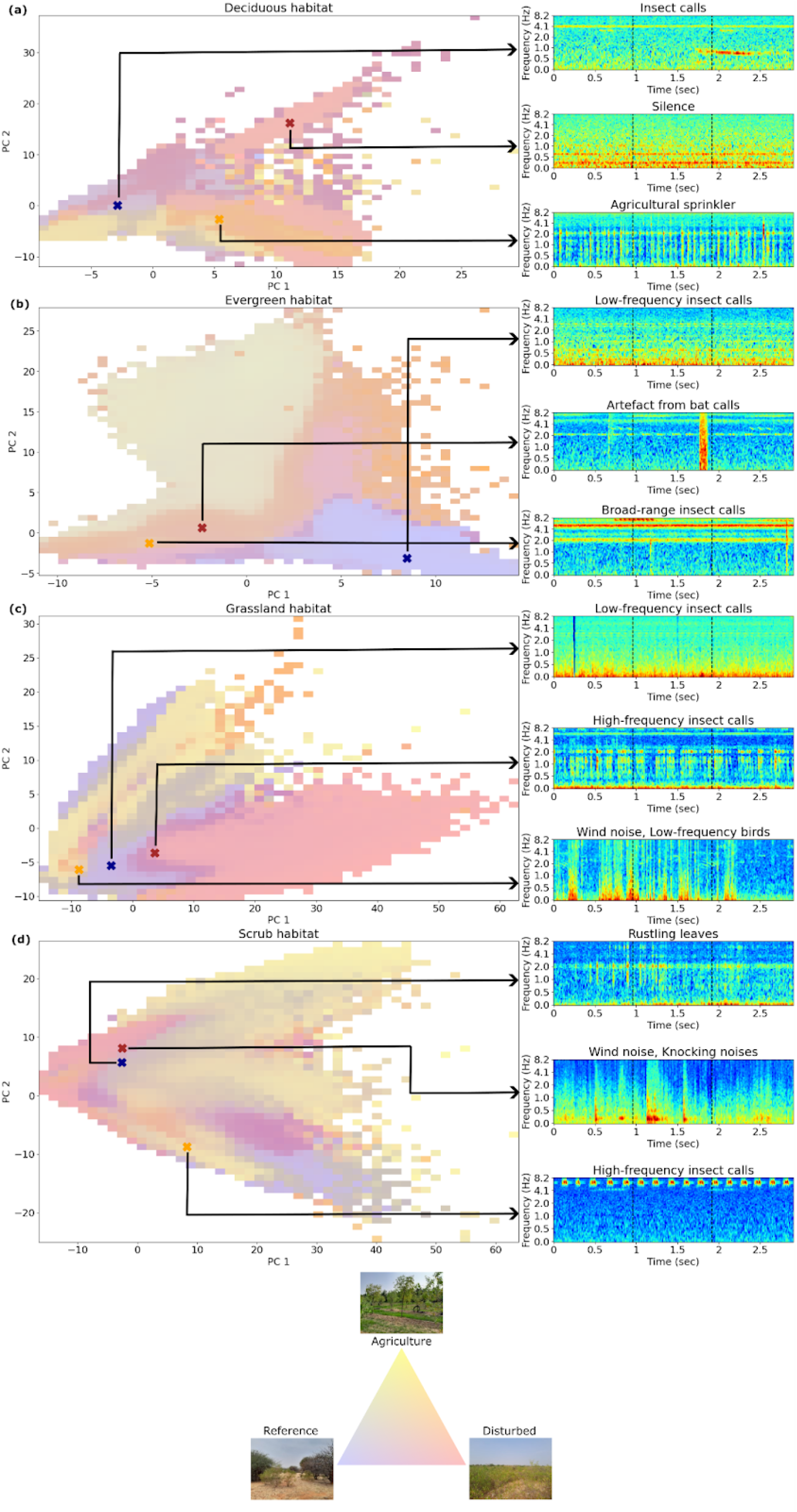
Identifying soundscape components relating to different land uses across habitats. The left-hand side plots show the spatial structure of acoustic embeddings of different land uses across habitats for the optimally thresholded model. The right-hand side shows example spectrograms sampled from GMM components of the three land uses based on the proportion of true positives. Insect sounds are the most predictive soundscape elements for land uses.

### Investigating GMM components gives biological insights into model predictions

We analysed and labelled audio samples from components of the aggregated habitat GMM classification models. We quantified the proportion of various soundscape elements in these components (Table 2) and ascertained component locations in the acoustic embedding space (Figure 5). Deciduous forest habitat provides the most ecologically comprehensible example of how the model made land use predictions. Most of the sampled audio in reference land use contained insect sounds, while agricultural land use exclusively contained noise from agricultural sprinklers, highlighting the distinguishing features for both land uses in the habitat and showcasing the importance of anthropophony in estimating habitat quality, especially in highly anthropogenically active land uses. For the evergreen forest habitat, all three land uses contained insect sounds in a very high proportion. However, the frequency bands these insects called in varied across land use, with insects in the reference land use calling around 2.5 kHz and 3 kHz, while insects in agriculture land use called at higher frequencies.

**Table 2.**
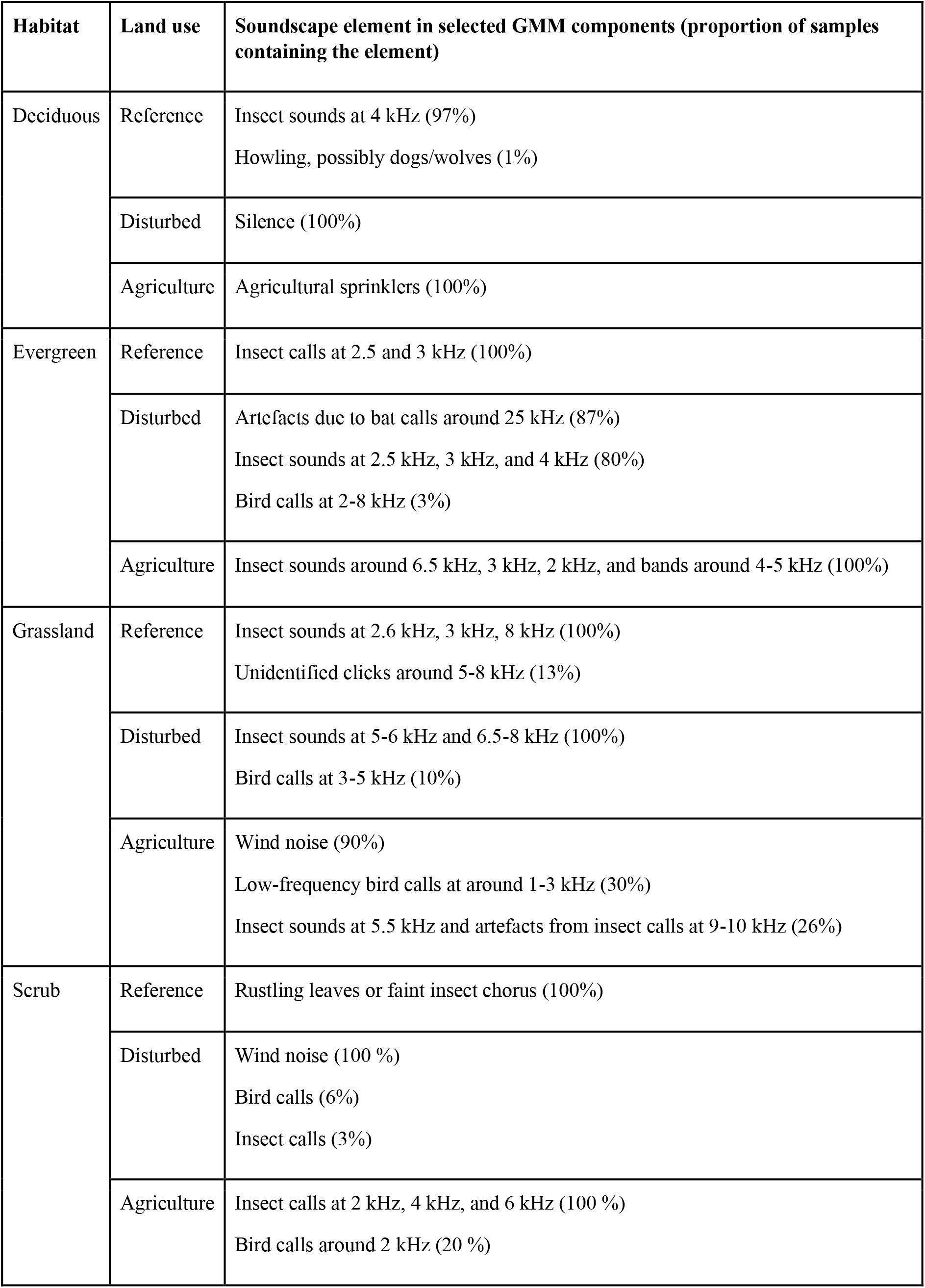
Proportion of soundscape elements in audio sampled from GMM components across habitats.

Similarly, for grassland habitat, audio samples from reference and disturbed land use almost always contained insect vocalisations but at different frequency bands, with insects in disturbed land uses not calling at lower frequencies. The most important distinguishing factor in grassland agriculture land use was wind noise, followed by low-frequency bird calls. The scrub habitat was also dominated by wind, with all sampled recordings from disturbed land use containing high wind noise and reference land use recordings containing noises from rustling leaves. Audio from agricultural land use was more silent, composed chiefly of insect vocalisations at 2, 4, and 6 kHz and bird calls around 2 kHz. Some loud insect vocalisations at around 9-10 kHz led to artefacts in agricultural land use in grassland in around 26% of the sampled recordings. Artifacts from bat vocalisations at 25 kHz were the most predictive soundscape elements for disturbed land use in evergreen forests. A detailed list of the various soundscape elements and their occurrence in GMM component samples is given in Table 2.

These soundscape elements were also well-segregated in the acoustic embedding space. For example, agricultural sprinkler noise, an indicator of agricultural land use in deciduous habitat, falls in the yellow region of the deciduous acoustic embedding space, restricting its presence only in agricultural land use (Figure 5a). Insects vocalising at around 6.5 kHz in grasslands were primarily present in disturbed land use (Figure 5c). However, similar calls in the evergreen habitat were present in overlapping regions of disturbed and agricultural land use, indicating the presence of such insects in both land uses (Figure 5b). The most informative components contained biophonic sounds, such as insect, bat, or bird vocalisations, depicting the ecological grounding of our model inferences (Figure 5).

## Discussion

We have demonstrated a novel framework for building an interpretable classifier model that can also help sort out uninformative acoustic data from potentially informative audio data using GMMs and VGGish acoustic feature embeddings. Using an example of land use classification across different habitats in tropical India, we demonstrated how to use the GMMs to make a classification model and threshold predictions to drop uninformative data in a generalised framework. Further, we showed that the framework could be used as a data exploration tool to extract important soundscape components and enquire about the inner workings of the model, making it more interpretable and helping ensure that the classifications made by the model have some ecological basis.

Although our GMM classifier model performed well across most habitats, the model performance in scrub habitat was underwhelming. This same pattern of model performance was observed for other classification models as well (Table 1), which might have arisen from methodological or ecological reasons. The major methodological issues for scrub habitat might be site selection and our method of choosing optimal thresholds. The plots selected as reference and disturbed for scrub habitat were more similar to each other than plots selected as reference and disturbed in other habitats, due to pervasive impacts of people across the wider scrub landscape. Replicating the study in another region with less-disturbed reference scrublands might be expected to yield clearer results. Moreover, any slight differences in soundscapes that may exist may have been masked by the optimal threshold selection we implemented. Since we picked a threshold based on the first inflection point rather than a hard threshold based on a cut-off value for precision, the model might have retained have uninformative data that needs to be dropped. Alternatively, the weak performance of the model in scrub habitats might also be ecological, in that there just may not be any discriminating information in the soundscapes of the reference and disturbed habitats: they might just sound the same.

The feature extractor we used to derive acoustic feature embeddings was VGGish, trained on Google’s AudioSet, a labelled sound library curated from millions of YouTube videos (Gemmeke et al., 2017). One of the core issues about the use of such feature extractor models is probably the bias in the training data; since the model was not initially intended to be used for land use classification, the embeddings it produces might not be relevant for the task (Gibb et al., 2024). However, we chose VGGish since it is trained on a broad dataset and can be used to identify broad soundscape components that can help estimate habitat quality (Sethi et al., 2020). Issues have also been raised regarding whether these habitat quality classifications use ecologically relevant soundscape markers (Gibb et al., 2024). Our results show that the VGGish model does pick out biophony across habitats, and these, in turn, help the model make land use classifications. One might argue that most of the biophony picked up by the model is insect sounds, indicating that VGGish might not be the best choice for other taxa. However, we would like to reiterate that there are vocalisations of other animals, such as some artefacts from bat calls and bird vocalisations. Previous studies have also shown that insect vocalisations are critical for acoustic habitat quality estimation (Gibb et al., 2024; Rappaport et al., 2022). Thus, the best-selected components we analysed may mostly contain insect vocalisations because they are more informative in distinguishing different land uses than vocalisations of other animals.

An issue with this feature extractor and most feature extractors in use for ecoacoustic classifications is the frequency range in which these CNNs function; most of them work within the 16 kHz range (Gibb et al., 2024; Kahl et al., 2021; Sethi et al., 2020). This omits important biophonic sounds such as bat vocalisations and other insect sounds (e.g. katydids) that might be relevant to assessing habitat quality (De Conno et al., 2018; Diwakar & Balakrishnan, 2012). Future work might include trying out multiple feature extractors, comparing their sensitivities to different sound classes, and determining if we can generate diverse information using complementary feature extractors.

Advances in interpretable machine learning models have primarily been geared towards devising better feature extraction tools (Eldridge et al., 2018; Gibb et al., 2024; Nieto-Mora et al., 2024; Sethi et al., 2020). Some feature extractors are better at certain specific tasks, e.g., BirdNet for species classification or VGGish and VAEs for habitat classification (Gibb et al., 2024; McGinn et al., 2023; Nieto-Mora et al., 2024; Sethi et al., 2020). Since our framework is modular with respect to the feature extractor used, one can use our framework as a filter to try out various feature extractors and select the best-performing feature extraction pipeline for a given task. Unlike current advances in the field that aim to build newer feature extraction tools or automated models, our framework can also be applied to previous classification studies using any black box feature extractor pipeline to identify soundscape components relevant to the classification tasks and investigate the ecological relevance of various identified components. Along with interpretability, the incorporation of task-driven information filtering makes our framework a much more powerful tool for ecoacoustic data analysis, particularly for building trustworthy and performant classifiers.

The methods developed here can be applied to a broad range of tasks in ecology and conservation. Our method provides a way to estimate the acoustic structure of a soundscape with a feature extractor. Monitoring changes in acoustic structure and soundscape across multiple seasons can be helpful for long-term habitat monitoring and could lead to the characterisation of ecological restoration and ecoacoustic succession, which can be defined as changes in soundscape structure across time as a landscape goes through ecological succession (Borker et al., 2020; Meyer et al., 2022; Rappaport et al., 2022). Combined with the occurrence data of various species, soundscape structure can also be used to identify and quantify ecologically relevant soundscape components correlating to habitat suitability (Sánchez-Giraldo et al., 2021). Thus, it provides a method to derive biologically meaningful interpretations of habitat suitability studies based on soundscapes (Sethi et al., 2022).

Our framework also provides methods to isolate and identify important soundscape elements for a classification model. This can help probe into classification models’ inner functioning and ascertain the trustworthiness of predictions (Gibb et al., 2024; Ribeiro et al., 2016). It can also provide insight into task-specific soundscape elements that aid in acoustic classification (Gibb et al., 2024; Nieto-Mora et al., 2024). Finally, the process of thresholding demonstrated in our study can be an essential tool to sort informative audio recordings from large amounts of audio recordings, especially in long-term, large-scale monitoring projects, explicitly aiding in dataset curation and decreasing model bias (Gamillo, 2023; Roe et al., 2021; Sethi et al., 2021).

## Conclusion

We have addressed two pressing issues in ecoacoustics and passive acoustic monitoring: the interpretability of models and the filtering of sparse information from large volumes of soundscape data. We present a framework that can be built upon any feature extraction pipeline. The framework can be used as a data exploration and filtering tool to improve classification and help curate training data for classification tasks. Our approach can also be used to understand the underlying reasons for decisions taken by a classifier while using certain soundscape features, thus inducting interpretability into an otherwise black box representation and method of classification.

## Data Availability

Code to reproduce figures and results presented in this manuscript can be found at https://doi.org/10.5281/zenodo.14045210, with the data required to run the code hosted at https://zenodo.org/records/13772137.

## Acknowledgements

We thank Prof. Mahesh Sankaran for allowing use of his field facilities, Mr. P. R. Bhat and Deepak Shetty for helping with site selection and the Karnataka State Forest Department for permitting us to sample in Tograhalli Reserved Forest. We thank Prof. D. N. Rao for facilitating fieldwork at the Challakere campus of IISc. We also thank the owners of private agricultural lands and estates for permitting us to sample on their properties. We extend our gratitude to Deepak Shetty, Raghavendra, Kallesh, and Harshavardhan for all their hard work in the field during data collection. Thanks to all the folks of the RB Lab at CES and RE Lab at Imperial College for discussions and valuable feedback at various stages of the project. Part of the initial calculations were done using the Param Pravega supercomputer at SERC IISc, we thank the IISc Param Pravega team for their help and support during running those computations.

## Author Contributions

All authors jointly contributed to the conception and design of the study. AO conducted the field surveys and cleaned up the data. AO and SS jointly designed the data analysis approaches, with regular input from RB and RE. AO and SS led manuscript writing; all authors contributed critically to the drafts and gave final approval for publication.

