## Supplemental Tables and Figures for "Interpretable and Robust Machine Learning for Exploring and Classifying Soundscape Data"

### Supplementary Information

**Table T1: Details of sampling sites.** The table below details the various habitats, climatic, and anthropogenic features of the different land use type plots set up in each sampling site

| **Sampling Site** | **Sampling Site Location** | **MAT*(°C)** | **MATR* (°C)** | **MAP* (mm)** | **Habitat type** | **High-quality plots** | **Medium quality plots** | **Low-quality plots** | **Hours of recorded audio** | **Final no. of plots sampled+** |
|---|---|---|---|---|---|---|---|---|---|---|
| Mulgund | 20 km Southwest of Sirsi town | 23.8 | 15.5 | 4166.25 | Evergreen forest | Reserve forest area with minimal anthropogenic activity | Unprotected forests surrounding areca plantation where the farmers collect leaves and branches for mulching | Areca and banana plantations | 850.96 | H = 4<br>M = 4<br>L = 4 |
| Tograhalli | 20 km Northeast of Siri town | 23.5 | 16.8 | 2008 | Deciduous forest | Reserve forest area with minimal anthropogenic activity | Teak plantations are regularly monitored by the government | Areca plantations | 796.12 | H = 5<br>M = 5<br>L = 4 |
| Khudapura | 15 km Northwest of Challakere town | 25.8 | 19.8 | 457.7 | Thorny and Scrub forest | Fenced scrubland inside the IISc Challakere campus | Unfenced scrublands outside the IISc campus with intensive grazing | Coconut plantations | 743.06 | H = 4<br>M = 4<br>L = 5 |
| Chetnal | 25 km North of the town of Bidar | 26.4 | 27.4 | 979.4 | Grassland | A large patch of grassland with minimal grazing | Fragmented grasslands near agricultural fields used for grazing | Sorghum and Safflower plantations | 786.24 | H = 5<br>M = 4<br>L = 5 |

* MAT = Mean Annual Temperature; MATR = Mean Annual Temperature Range; MAP = Mean Annual Precipitation. + In some sites, the deployed recorder malfunctioned; thus, the number of final plots sampled is slightly less than the number of plots where recorders were put up.

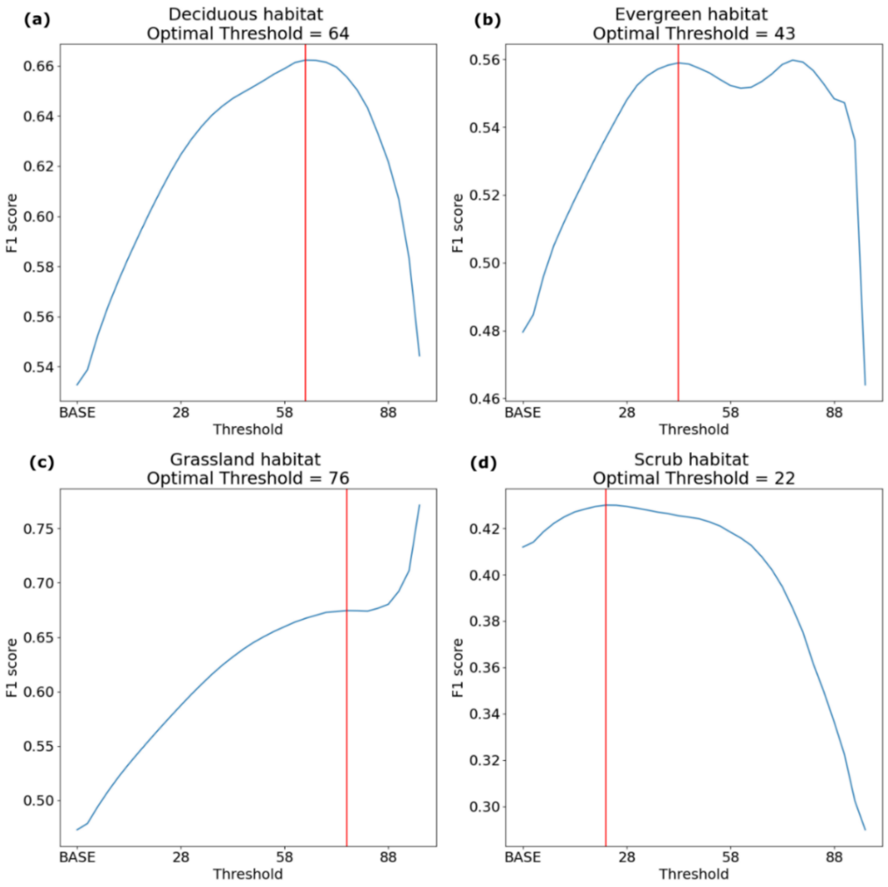


**Figure S1: F1 score vs threshold plot**.

The plots above show the change in model F1 score as we increase the threshold. The optimal threshold was chosen as the first inflexion point of the curve in each habitat (indicated by the red vertical line).

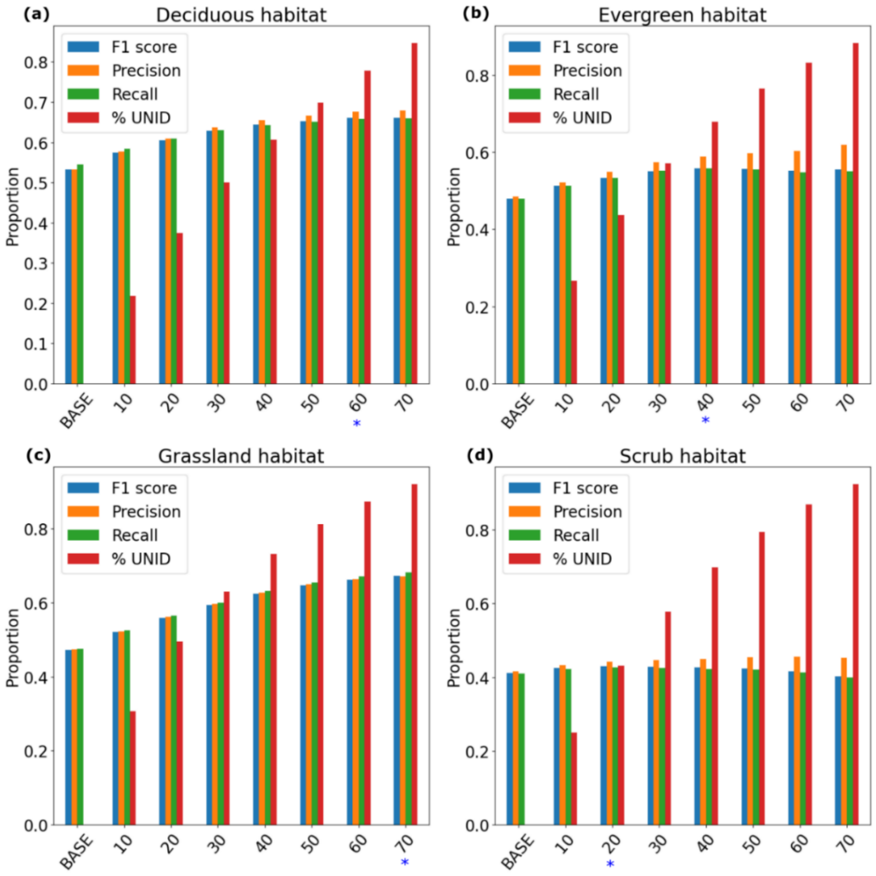


**Figure S2: Model accuracy metrics vs threshold.**

Model accuracy (precision, recall, F1 score) increases with increasing threshold value across habitats. However, in scrub habitat, the model F1 score and recall decrease slightly after a threshold of 20 percentile. Blue stars represent threshold values closest to the optimal threshold for each habitat.
